# Differential effects of ethanol on behavior and GABA_A_ receptor subunit expression in zebrafish (*Danio rerio*) with alternative stress coping styles

**DOI:** 10.1101/829515

**Authors:** Alexander C. Goodman, Ryan Y. Wong

## Abstract

Variation in stress responses between individuals is linked to factors ranging from stress coping styles to sensitivity of neurotransmitter systems. Many anxiolytic compounds (e.g. ethanol) can increase stressor engagement through modulation of neurotransmitter systems and are used to investigate stress response mechanisms. Here we assessed the role of the GABA_A_ system on the variation of the behavioral stress response by comparing individuals differing in stress coping styles that were chronically treated with ethanol. Specifically, we investigated resulting changes in stress-related behavior and whole-brain GABA_A_ receptor subunits (*gabra1, gabra2, gabrd*, & *gabrg2*) in response to a novelty stressor. There were significant main and interaction effects on two stress-related behaviors, where the ethanol-treated proactive individuals showed lower stress-related behaviors than their reactive counterparts. Proactive individuals showed significantly higher expression of *gabra1, gabra2*, and *gabrg2* compared to reactive individuals and ethanol treatment resulted in upregulation of *gabra1* and *gabrg2* in both stress coping styles. These results show that differences in stress-related behaviors between stress coping styles may be facilitated in part by expression of select GABA_A_ receptor subunits.

## Introduction

While an organism’s stress response is essential to its survival, not all conspecifics exhibit similar responses and often differ both behaviorally and physiologically^1–5^. Across many taxa there exists two alternative correlated suites of behavioral and physiological responses to stressors known as the proactive and reactive stress coping styles^2,3,5–7^. Proactive individuals actively engage stressors and characteristically exhibit a lower whole-body cortisol response compared to reactive individuals in response to novelty^2,3,5,8–10^. Additionally, proactive and reactive individuals differ in expression of key neurotransmitter receptors related to stress and anxiety, such as serotonin, dopamine, and GABA (γ-amino butyric acid) receptors^2,3,11,12^. Drugs designed to target such systems are often employed to study a neurotransmitter’s influence on stress-related behaviors^13–15^. Therefore, pharmaceuticals can be used to investigate underlying differences in the molecular mechanisms between stress coping styles.

Dysregulation of the GABAergic, serotoninergic, and the glutamatergic systems often contribute to a disproportional behavioral stress response^13,16^, which, if sustained over an extended period of time, can be classified as an anxiety disorder^17,18^. GABAergic system dysfunction is thought to contribute to the underlying etiology of anxiety-related disorders^19,20^. GABA_A_ receptor (GABA_A_R) agonists, such as ethanol, allow for positive modulation of the GABAergic system to produce an anxiolytic response, while antagonists result in an anxiogenic response^13,16,21–28^. GABA-acting drugs influence the expression of the protein subunits that make up the receptor subtype as well^29,30^. For example, rodents exposed to GABAA agonists show an increase in expression of the α_1_-, α_2_-, and δ-subunits of the GABA_A_R, while expression of the γ_2_-subunit decreases^31–34^. Studies utilizing zebrafish similarly show that ethanol administration produces anxiolytic behavioral effects^13,23,24,26,35,36^. While there are baseline differences in mRNA expression of both GABA_A_ and GABA_B_ receptors between zebrafish with the proactive or reactive stress coping style^12^, how these drugs differentially influence both the behavior and GABAergic response while taking into account an individual’s stress coping style is not understood.

Zebrafish (*Danio rerio*) is a widely used model to understand the effects of pharmaceuticals on stress and anxiety-related behaviors and physiology due in part to their conserved behavioral, neuroanatomical, pharmacological and transcriptional stress responses with mammals and other species^13–15,24,37–41^. Furthermore, wild and laboratory strains of zebrafish show the proactive and reactive stress coping styles^5,6^. These coping styles in zebrafish display differences in genetic backgrounds, behavior and neuroendocrine responses to stressors that are consistent with what has been documented in birds and mammals^42–44^. Only recently are studies beginning to demonstrate the roles of synaptic plasticity and neurotransmitter system regulation in facilitating the display of alternative stress coping styles in zebrafish^5,7,12,45–47^. Hence zebrafish can serve as a useful system to study the neuromolecular variations between stress coping styles through the use of GABA-acting drugs.

In this study, we assessed the effects of ethanol treatment on stress-related behavior and GABA_A_R subunit gene expression in two zebrafish lines selectively bred to display the proactive and reactive stress coping styles. Specifically, we quantified expression of four genes encoding for the α_1_-, α_2_-, δ-, and γ_2_-subunits of the GABA_A_R (*gabra1, gabra2, gadrd*, and *gabrg2*, respectively; ^48^. We hypothesized that ethanol treatment will reduce stress-related behaviors (e.g. exploratory behavior) in both lines of zebrafish with a greater anxiolytic response for the reactive line. Additionally, based on previous literature we predicted to see an increase in mRNA expression of α_1_-, α_2_-, δ-subunits and decrease expression of the γ_2_-subunit for both lines but the magnitude of the effect would be greater in the reactive line^31–34^. Understanding how a GABA_A_R agonist impacts GABA neurotransmission between the two coping styles will give insight into one mechanism that may explain differences in their stress and anxiety-related behavioral responses.

## Materials and Methods

### Subjects

In this study, we used the high-stationary behavior (HSB) and low-stationary behavior (LSB) lines of zebrafish (*Danio rerio*). These two lines exhibit differences in stress-related behaviors across multiple behavioral assays, learning and memory, glucocorticoid responses, neurotranscriptome profiles, and morphology consistent with the reactive and proactive stress coping styles^5,6,10,12,45,47,49,50^. Therefore, we consider any fish from the HSB or LSB lines to have the reactive or proactive stress coping style, respectively. Lines were generated starting from a wild-caught population from Gaighata in West Bengal, India and are maintained through a bidirectional selective breeding paradigm on behavioral stress response to a novelty stressor ^5^. Both lines were 12 to 15 months post-fertilization when testing began and underwent 11 generations of selective breeding. Prior to testing, fish were housed in 40-liter mixed-sex tanks on a recirculating system. Water temperature was set at 27°C. Fish were kept on a 14:10 L/D cycle and fed twice daily with Tetramin Tropical Flakes (Tetra, USA). All procedures and experiments were approved by the Institutional Animal Care and Use Committee of the University of Nebraska at Omaha/University of Nebraska Medical Center (17-070-09-FC).

### Pharmacological manipulation

To identify a biologically relevant ethanol dose, we conducted a pilot dose-response study. We chronically administered ethanol of varying concentrations and durations to both lines followed by a behavioral stress assay (Novel Tank Diving Test) to measure anxiety-related behaviors (see below). Ethanol treatment began at 0.25% v/v over a period of seven days. Concentration and duration were progressively increased until an anxiolytic effect was observed in both lines of zebrafish without drug-impaired locomotion (i.e. significant change in depth preference with no significant difference or decrease in distance traveled and stationary time relative to control fish). We used total distance traveled and total stationary time during the trials as proxies for locomotion to ensure the chosen concentration of ethanol was not impairing the fish’s ability to swim. We tested treatment durations from 7 days (0.25%, 0.4%, 0.5%, 0.75%, 1%, 1.15%, 1.25%, and 1.5% ethanol), 10 days (0.5% ethanol), up to 14 days (0.5% and 0.75% ethanol) (Figure S1, Tables S1-S4). There were significant main effects of ethanol concentration on time spent in the top half of the tank for both the HSB and LSB lines at the 14-day duration (HSB: *χ*^*2*^(2) = 19.293, *p* ≤ 0.001; LSB: *χ*^*2*^(2) = 11.330, *p* ≤ 0.01). Post-hoc analysis revealed fish treated with 0.75% ethanol concentration showed an increase in time spent in the top half of the tank compared to 0.0% concentration for both the HSB and LSB line (HSB: *U* = 18.000, *p* ≤ 0.001; LSB: *U* = 49.500, *p* ≤ 0.001; Table S4) with no drug-impaired locomotion. Therefore, we selected the 0.75% ethanol for two weeks treatment regime for this study.

Using a modified protocol for chronic ethanol administration in zebrafish^24^, groups of six fish were housed in a 3-liter trapezoidal tank (15.2 height × 27.9 top × 22.5 bottom × 11.4 cm width; Pentair Aquatic Ecosystems) throughout the treatment period. The tank contained either 2-liters of 0.75% ethanol (v/v; Sigma-Aldrich) or 2-liters of system water as a control over the span of 14 days. Every two days we replaced the entire water in each tank with fresh ethanol or system water. At the end of 14 days, a group of fish was used for either behavioral testing or for quantification of whole-brain GABA_A_R subunit mRNA expression. We randomly selected 36 individuals from each of the HSB and LSB lines to be behaviorally tested (*N* = 18 for each treatment group). We used a different set of 36 individuals from each line (*N* =18 for each treatment group) for quantification of GABA_A_R subunit expression. Some fish were lost during the treatment period resulting in final sample sizes of 32 individuals from the HSB (*N* = 15 treated, 17 control; *Female* = 13, *Male* = 19) line and 33 from the LSB (*N* = 16 treated, 17 control; *Female* = 14, *Male* = 19) that were behaviorally tested using the NTDT. A total of 34 individuals from the HSB (*N* = 17 treated, 17 control; *Female* = 15, *Male* = 19) line and 35 from the LSB (*N* = 17 treated, 18 control; *Female* = 18, *Male* = 17) were used for GABA_A_R subunit quantification.

### Behavioral Testing

Following the 14^th^ day of treatment, fish were exposed to a novelty stressor by placing them into the Novel Tank Diving Test (NTDT) assay following established procedures^5,10,49^. Reduced transitions to and time spent in the top half of the tank are indicators of heightened stress and anxiety^5,24,51^. In brief, fish were netted from their treatment tanks and individually placed in a clear 3-liter trapezoidal tank (15.2 height × 27.9 top × 22.5 bottom × 11.4 cm width; Pentair Aquatic Ecosystems) filled with 2-liters of system water. We video-recorded the fish for six minutes and quantified behaviors using an automated tracking software (Noldus Ethovision XT, Wageningen, Netherlands) as previously described^6^. Specifically, we used the software to virtually partition the tank into top and bottom halves to measure the number of transitions to the top portion of the tank, time spent in the top portion of the tank (s), total distance traveled (cm), and stationary time (s). The subject was considered stationary if it was moving less than 0.5 cm/s. Stationary time and distance travels were used as proxies for locomotor activity to assess whether or not ethanol treatment impaired general locomotor activity. Testing occurred between 0800-1700 hours.

### Quantification of GABA_A_R subunit expression

We quantified whole-brain expression of four genes that encode for GABA_A_ receptor subunits (*gabra1, gabra2, gadrd*, and *gabrg2;* Table S5), and one housekeeping gene (*ef1a*) using quantitative reverse transcriptase PCR (qRT-PCR) following established protocols^12,49,50^. In brief, whole brains were homogenized with 50-100 µL of zirconium oxide beads (Bullet Blender, Next Advanced) in Tri Reagent (Sigma-Aldrich). Then, we extracted RNA and removed genomic DNA using column filtration (PureLink RNA Mini Kit, Ambion). We subsequently synthesized cDNA using both random hexamers and oligo(dT)_20_ primers. (SuperScript IV First-Strand Synthesis System for qRT-PCR (Invitrogen). Finally, we purified the cDNA using Amicon Ultracentrifugal filters (Millipore). We carried out all protocols according to each manufacturers’ protocol.

We ran the qRT-PCR on QuantStudio 7 Flex Real-Time PCR System (Applied Biosystems) using SYBR green detection chemistry (PowerUp SYBR Green Master Mix, Applied Biosystems). The primers were designed using Primer-Blast ^52^ with chosen primers either spanning exon-exon junctions or with the amplicon spanning exons where the intron region was over one kilobase (Table S5). Primer concentrations were 5 pmol for all genes. Reaction parameters for all genes were as follows: 2 minutes at 50°C, 2 minutes at 95°C, followed by 40 cycles at 95°C for 15 seconds then 60°C for 1 minute. We ran each sample in triplicate. We quantified expression using the relative standard curve method and normalized expression to an endogenous reference gene (*ef1a*). *ef1a* expression is stable across sex, tissue types, age, and chemical treatment in zebrafish^53^.

### Statistical Analysis

We used a generalized linear model (GLZ) in SPSS (Version 24) to assess changes in behaviors and gene expression because the data was not normally distributed. Line (HSB, LSB), sex (male, female) and treatment group (0.75% ethanol, control) were used as between-subject variables. As the relationship between body size and locomotion is well documented ^45,54–56^, we included standard length as a covariate. Since we did not find a significant main effect of sex on behavior (top transitions: *χ*^*2*^(1) = 2.385, *p* = 0.123; top time: *χ*^*2*^(1) = 0.852, *p* = 0.356; distance: *χ*^*2*^(1) = 0.682, *p* = 0.409; and stationary time: *χ*^*2*^(1) = 0.092, *p* = 0.762) or gene expression (*gabra1: χ*^*2*^(1) =0.036, *p* = 0.850; *gabra2: χ*^*2*^(1) = 0.382, *p* = 0.536; *gadrd: χ*^*2*^(1) = 1.942, *p* = 0.163; *gabrg2: χ*^*2*^(1) = 1.426, *p* = 0.232), we removed that variable from the analyses and used a simpler GLZ with line and treatment group as the only between-subject variables. For the post-hoc comparisons, we ran Mann-Whitney U tests and applied a Benjamini-Hochberg correction to correct for multiple comparisons ^57^. As our pilot and multiple other studies show that ethanol results in the decrease of stress and anxiety-related behavior^13,16,21–26^, we assessed significant differences in post-hoc comparisons of stress-related behaviors between treatment and control groups using one-tailed p-values. Significance of all other post-hoc comparisons used two-tailed p-values.

## Results

### Greater anxiolytic effect of ethanol on behavior in the LSB line

There were significant main effects of line on both top transitions (*χ*^*2*^(1) = 12.579, *p* ≤ 0.001) and time spent in the top half of the tank (*χ*^*2*^(1) = 10.215, *p* ≤ 0.001). LSB fish transitioned to (*U* = 281.500, *p* ≤ 0.001; Figure 1a) and spent significantly more time in the top half of the tank (*U* = 297.000, *p* ≤ 0.01; Figure 1b) than HSB fish. There were also significant main effects of treatment on both top transitions (*χ*^*2*^(1) = 28.054, *p* ≤ 0.001) and time spent in the top half of the tank (*χ*^*2*^(1) = 32.659, *p* ≤ 0.001). Ethanol-treated fish transitioned to (*U* = 234.500, *p*_*one-tail*_ ≤ 0.001) and spent significantly more time in the top half of the tank (*U* = 236.000, *p*_*one-tail*_ ≤ 0.001) than control fish. There was a significant line by treatment interaction effect for transitions to the top half of the tank (*χ*^*2*^(1) = 6.788, *p* ≤ 0.001) and time spent in the top half of the tank (*χ*^*2*^(1) = 8.182, *p* ≤ 0.01). Ethanol-treated LSB fish exhibited the most top transitions compared to the control HSB (*U* = 11.500, *p* ≤ 0.001), control LSB (*U* = 28.000, *p*_*one-tail*_ ≤ 0.001), and ethanol-treated HSB fish (*U* = 48.500, *p* ≤ 0.01). This pattern was also found for time spent in the top half with ethanol-treated LSB exhibiting the most time spent in the top half of the tank compared to control HSB (*U* = 15.000, *p* ≤ 0.01), control LSB (*U* = 27.500, *p*_*one-tail*_ ≤ 0.001), and ethanol-treated HSB fish (*U* = 49.000, *p* ≤ 0.01).

**Figure 1.**
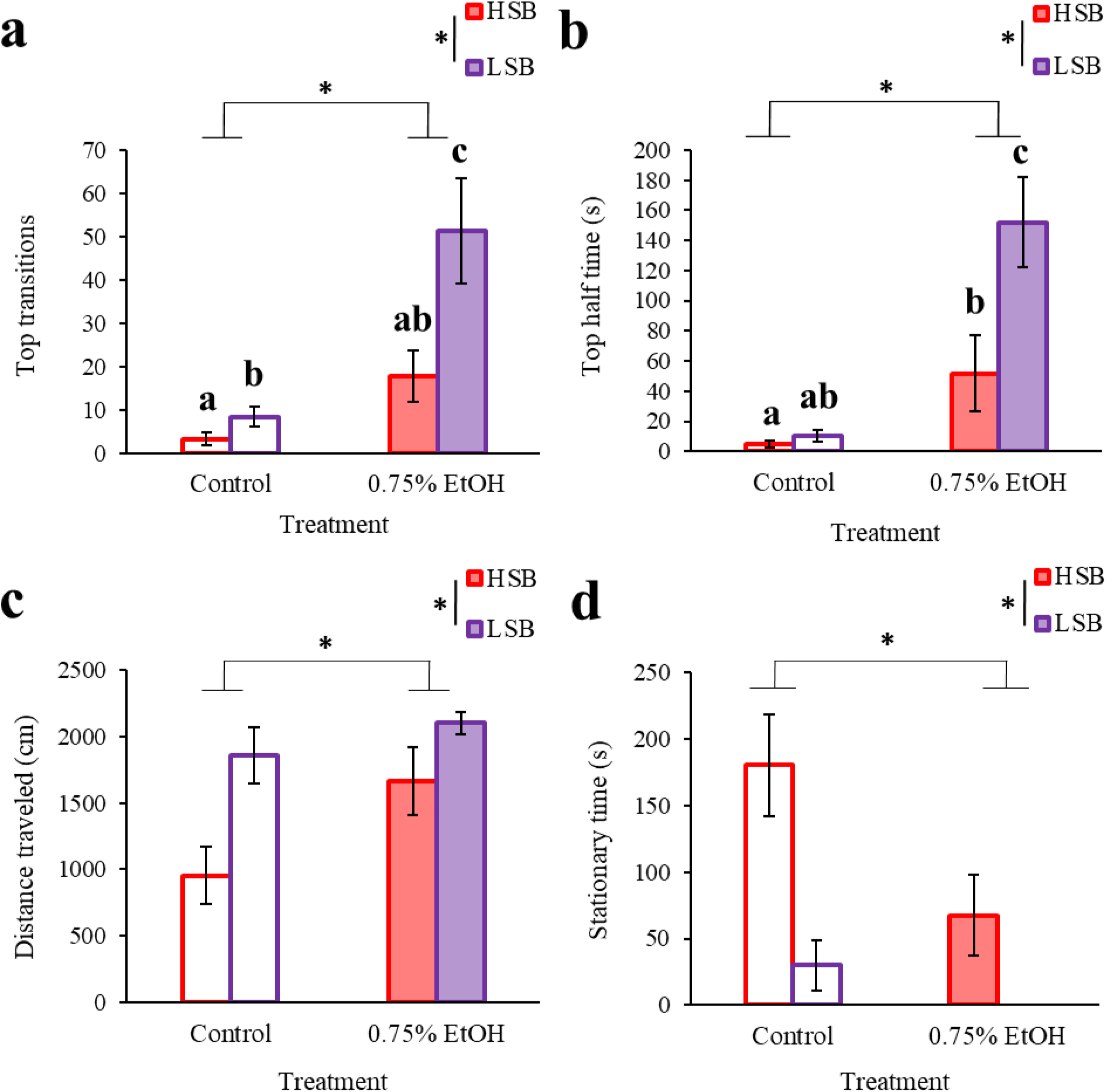
Differentiated ethanol treatment effect on anxiety-related behaviors between lines with no effect on locomotion. We measured top transitions (**a**), time in top half of the tank (**b**), distance traveled (**c**), and stationary time (**d**) for each treatment group. Control groups are represented by unfilled in bars, while ethanol-treated groups are represented by filled bars. HSB and LSB are red and purple, respectively. Data shown are mean ± 1 SEM. Significant line and treatment differences are indicated by an asterisk (*p* ≤ 0.05), while differences between groups are indicated by different lower case letters.

### No impaired locomotion from ethanol-treatment for both lines

There were significant line effects for total distance swam (*χ*^*2*^(1) = 11.378, *p* ≤ 0.001) and stationary time (*χ*^*2*^(1) = 18.173, *p* ≤ 0.001). LSB fish swam a significantly farther distance (*U* = 280.000; *p* ≤ 0.001; Figure 1c) and spent significantly less time stationary (*U* = 216.000; *p* ≤ 0.001; Figure 1d) than HSB fish. We also found significant treatment effects for total distance swam (*χ*^*2*^(1) = 5.729, *p* ≤ 0.05) and stationary time (*χ*^*2*^(1) = 7.831, *p* ≤ 0.01). Ethanol-treated fish traveled farther (*U* = 360.000; *p* ≤ 0.05) and spent less time stationary (*U* = 364.000; *p* ≤ 0.05) than control fish. There were not any significant line by treatment interaction effects for total distance travelled (*χ*^*2*^(1) = 1.391, *p* = 0.238) or stationary time (*χ*^*2*^(1) = 2.639, *p* = 0.104).

### Ethanol-treatment increases expression of α_1_ and γ_2_ GABA_A_R subunits

We found significant main effects of line on expression of *gabra1* (*χ*^*2*^(1) = 7.310, *p* ≤ 0.01), *gabra2* (*χ*^*2*^(1) = 8.235, *p* ≤ 0.01), and *gabrg2* (*χ*^*2*^(1) = 5.929, *p* ≤ 0.05), but not *gabrd* (*χ*^*2*^(1) = 0.023, *p* = 0.880). The LSB fish showed higher expression of the *α*_1_-(*U* = 372.000; *p* ≤ 0.05), *α*_2_-(*U* = 393.000; *p* ≤ 0.05), and *γ*_2_-subunit (*U* = 365.000; *p* ≤ 0.05) than the HSB fish (Figure 2a, 2b, and 2d). There were significant main effects of treatment on expression of *gabra1* (*χ*^*2*^(1) = 6.507, *p* ≤ 0.05) and *gabrg2* (*χ*^*2*^(1) = 7.220, *p* ≤ 0.05) but not *gabra2* (*χ*^*2*^(1) = 0.648, *p* = 0.421) or *gabrd* (*χ*^*2*^(1) = 2.042, *p* = 0.153). Ethanol-treated fish showed greater expression of the *α*_1_-(*U* = 393.500; *p* ≤ 0.05) and *γ*_2_-subunit (*U* = 386.000; *p* ≤ 0.05) than control fish. There were no significant line by treatment interaction effects for any of the four subunits (*gabra1*: *χ*^*2*^(1)= 1.339, *p* = 0.247; *gabra2*: *χ*^*2*^(1) = 0.073, *p* = 0.787; *gabrd*: *χ*^*2*^(1)= 0.832, *p* = 0.362; *gabrg2*: *χ*^*2*^(1)= 0.659, *p* = 0.417).

**Figure 2.**
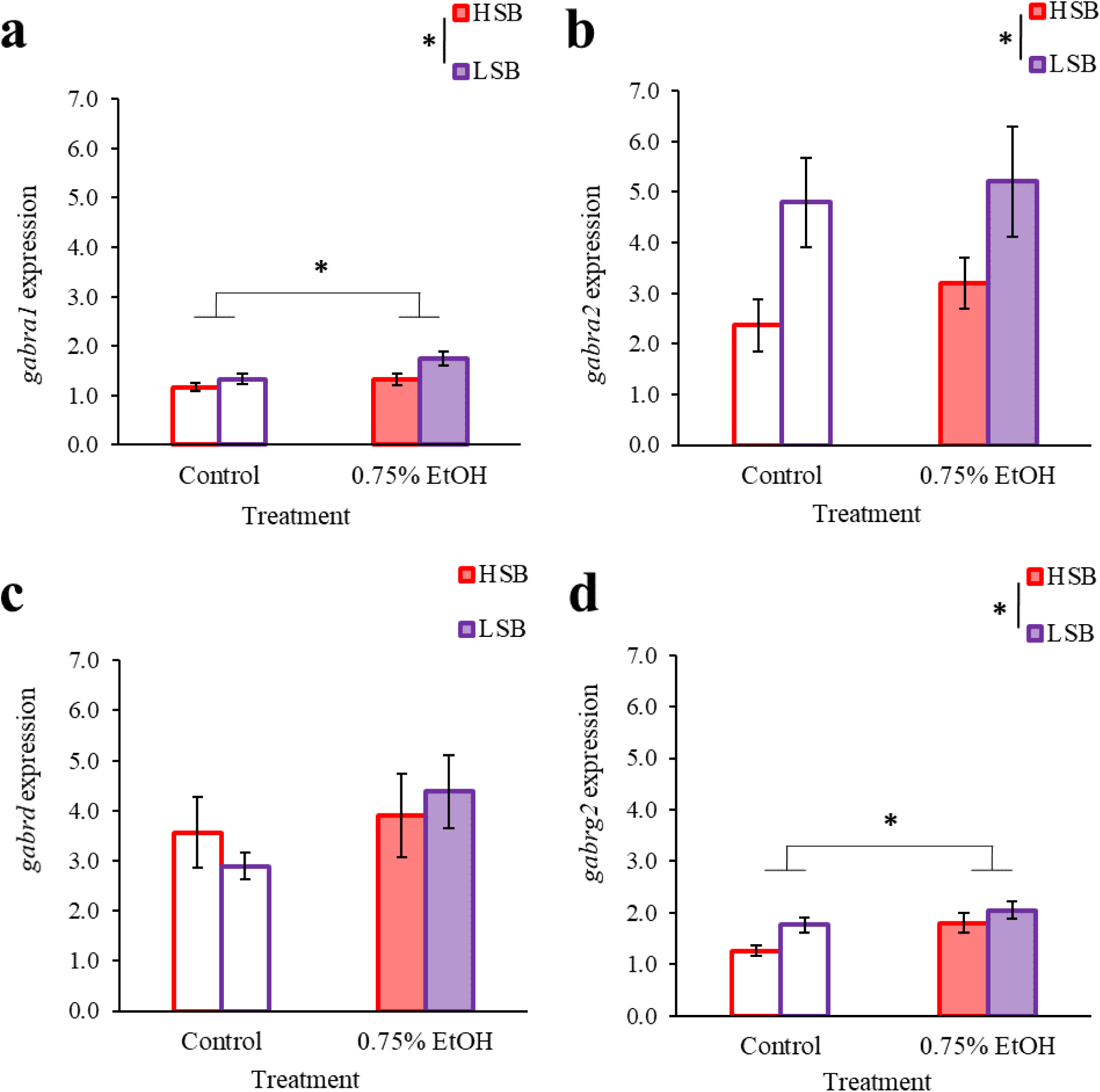
Effect of line and treatment on GABA_A_ receptor subunits. Normalized expression of *gabra1* (**a**), *gabra2* (**b**), *gabrd* (**c**), and *gabrg2* (**d**) for each treatment group following treatment. Control groups are represented by unfilled in bars, while ethanol-treated groups are represented by filled bars. HSB and LSB are red and purple, respectively. Data shown are mean ± 1 SEM. Significant differences are indicated by an asterisk (*p* ≤ 0.05).

## Discussion

GABA_A_ agonists, such as ethanol, produce an anxiolytic response across many taxa^13,16,21–26,58^. Through the use of these stress-reducing compounds, we can investigate the role of the GABAergic system in facilitating the expression of a stress coping style. In this study, we assessed both the behavioral and molecular responses of ethanol treatment between proactive (LSB) and reactive (HSB) lines of zebrafish. We found that while chronic ethanol treatment decreased stress-related behaviors in both lines, ethanol treatment had a greater anxiolytic effect on LSB line. The differences in stress-related behavior are linked to differential GABA_A_R receptor subunit expression between the lines (α_1_-, α_2_-, and γ_2_-subunits) or in response to ethanol treatment (α_1_-, and γ_2_-subunits). The results suggest molecular differences in the GABAergic neurotransmitter system contribute to the variation in stress-related behaviors between the two stress coping styles.

The anxiolytic behavioral response to ethanol in zebrafish is well documented^13,24,26,35,36^, but the effect of an individual’s stress coping style on the response to GABA_A_R agonists has only been recently investigated. We predicted that treatment with a GABA agonist would have a greater anxiolytic effect on both stress-related behaviors and GABA_A_R receptor subunit expression in the reactive stress coping style than the proactive stress coping style. As expected, we found that both the LSB (proactive) and HSB (reactive) lines of zebrafish displayed a decrease in anxiety-related behaviors following ethanol treatment. Surprisingly, the proactive individuals showed a greater anxiolytic response than the reactive individuals. To our knowledge, only one other study accounted for stress coping style when examining the anxiolytic effects of ethanol in zebrafish^59^. In that study, acute ethanol treatment resulted in a greater anxiolytic effect (fish spent more time in an area of the tank furthest from conspecifics) on reactive fish, while proactive fish increased their stress-related behaviors^59^. We speculate the opposing observations between our studies could be due to differences in treatment length (60 minutes vs. 2 weeks), social stress buffering (social vs. isolation), and assignment of stress coping style (behavioral screen vs. selectively bred lines). Regardless, ethanol is known to have an anxiolytic effect and the behavioral results from the prior and current studies suggest that an individual’s stress coping style can modulate the magnitude of the effect.

More generally, the line-specific responses to ethanol treatment we observed are consistent with other studies in zebrafish and rodents^60–65^. We found that the LSB line of zebrafish showed the greatest increase in transitions to and time spent in the top half of the tank during the NTDT compared to the HSB line. This line-specific response can be seen in other zebrafish studies. Laboratory lines of zebrafish require a higher concentration of ethanol to match exploratory behavior of wild-caught lines, while wild-caught lines exhibit abolishment of shoaling behavior at higher concentrations of ethanol^60–62^. Rodents selectively bred to exhibit diverging novelty-seeking behaviors show differing levels of responsiveness to ethanol^63–65^. Maintaining laboratory and selectively bred lines of animals simultaneously results in line-specific genetic backgrounds. For example, the HSB and LSB zebrafish lines used here show distinct whole-brain transcriptome profiles^12,50^ and the divergent novelty-seeking rodent lines differ in neuropeptide gene expression relating to the dopaminergic system^64,65^, suggesting that an individual’s behavioral response can be influenced by its genetic profile and underlying expression of neurotransmitters. Altogether our results show that differences in molecular mechanisms can contribute to the alternative behavioral stress-response between stress coping styles.

Unexpectedly, the proactive line (LSB) showed a greater anxiolytic behavioral response to ethanol than the reactive stress coping style line (HSB). It is possible that the higher expression of α_1_-, α_2_-, and γ_2_-subunits GABA_A_ receptor subunits we observed in this study in the proactive zebrafish facilitated a greater anxiolytic response to ethanol treatment. In rodents, removal of the α_2_-subunit results in the abolishment of the anxiolytic effect for both ethanol and other benzodiazepines^66,67^, suggesting this is a critical subunit needed for ethanol’s anxiolytic effect. We hypothesize that higher expression of these subunits in our proactive line may allow for greater sensitivity of GABA_A_ receptor ligands leading to a greater anxiolytic response.

In addition to being differentially expressed between the two lines, expression of the α_1_-, and γ_2_-subunits increased as a result of ethanol treatment. These results are consistent with previous studies in rodents where α_1_-subunit increased expression with ethanol treatment^31–34^. This suggests that ethanol-induced modulation of this subunit may be a conserved response across taxa. Prior studies examining the change in the γ_2_-subunit expression to ethanol treatment show conflicting information^68–70^. While our results are consistent with studies showing lower expression of this particular subunit decreases stress-related behaviors, other studies have shown increased expression similarly leading to a reduction in stress-related behaviors. It has been hypothesized that the γ_2_-subunit increases the overall responsiveness of the GABA neurotransmitter system^70,71^. Our results are consistent with this hypothesis as the proactive line showed higher expression of the γ_2_-subunit and had a greater change in the anxiolytic behavioral response from a GABA_A_ receptor agonist (ethanol). Interestingly, knockouts of either the α_1_- or γ_2_-subunits do not abolish ethanol’s anxiolytic effect. Both wild type and α_1_-subunit knockout rodents display an anxiolytic response to GABA_A_ receptor agonists, but rodents with the knockout display a greater decrease in anxiety-related behaviors, such as time spent in the open and number of open arm entries in the elevated plus maze^72–74^. Results of previous studies assessing γ_2_-subunit knockouts on stress-related behaviors are inconsistent. Some studies found partial knockout of this receptor subtype decreases exploratory behavior in an open field test (i.e. increasing anxiety)^68,69^, while a more recent study found complete knockout of the subunit in dopaminergic neurons increases exploratory behavior^70^. While removal of the α_1_- or γ_2_-subunits alter behavior in the rodent animal model, the anxiolytic effect of GABA_A_ agonist is still present regardless of the presence in the GABA_A_ receptor. This suggests that the α_1_- and γ_2_-subunits particular subunits are sufficient but not necessary for the anxiolytic response and their increased expression in the current study may have facilitated the reduction of stress-related behavioral displays in both lines.

Of note, we did not observe any significant line by treatment interaction effects on expression of any of the examined GABA_A_ receptor subunits. It is possible that by looking at whole-brain expression levels, we masked brain-region specific responses that may have shown interaction effects. As the GABAergic system can be differentially modulated depending on length (acute vs chronic) of ethanol exposure^58,62,75^, we also cannot rule out the possibility that our results may change with acute ethanol exposure. Another interpretation is that the GABAergic system does not play a significant role in the differentiated anxiolytic behavioral effects of chronic ethanol exposure between stress coping styles in zebrafish. Rather, the anxiolytic effects could be mediated by another neurotransmitter system such as the dopaminergic or serotoninergic system. Prior studies in fish and rodents have documented that administration of ethanol and other anxiolytic compounds alter several neurotransmitter systems in addition to the target system^49,76–81^. Of note, a prior study showed that the proactive (LSB) line showed higher baseline expression of the DRD2 receptor compared to the reactive (HSB) line^12^. Given this receptor’s role in ethanol-induced activation of the mesolimbic dopaminergic reward pathway of the brain and drug-seeking and novelty exploration behaviors^82–84^, we speculate that the differences in the magnitude of the anxiolytic effects of chronic ethanol on behavior between the two stress coping style lines involve the dopaminergic system. Future studies are needed to assess the extent of ethanol effects on neurotransmitter systems beyond the GABA_A_ system between the two stress coping styles.

## Conclusions

In this study, we showed significant main effects of line on anxiety-related behaviors and GABA_A_R subunit expressions where individuals with the proactive stress coping style (LSB line) had lower anxiety-related behaviors and higher expression of the α_1_, α_2_, and γ_2_-subunits relative to reactive (HSB) individuals. This demonstrates that variation in behavioral responses to a novelty stressor may be explained by differences in the GABAergic system (e.g. GABA_A_R subunit expression) between the two stress coping styles. Intriguingly we observed a significant line by ethanol treatment interaction effects on stress and anxiety-related behaviors. Chronic ethanol treatment had a surprisingly greater anxiolytic effect on proactive individuals, which suggests that ethanol alters the underlying neuromolecular mechanisms in a coping style-specific manner. However, the lack of an interaction effect between line and treatment on any of the four measured GABA_A_R subunits leads us to speculate that the differences in the magnitude of effect between the lines induced by chronic ethanol treatment may be mediated by a neurotransmitter system other than the GABAergic system. More broadly, this study shows that differences in stress and anxiety-related behaviors between the proactive and reactive stress coping styles are due in part to differences in the GABAergic system but any coping-style specific anxiolytic behavioral effects of chronic ethanol exposure likely involve other neurotransmitter systems.

## Acknowledgements

We are grateful to D. Revers, S. Roundtree, A. Park, and N. Mohamed for zebrafish husbandry. We thank M. Baker, K. Cullen, R. Patterson, and other members of the Wong lab for helpful discussions. This study was supported by the UNO Fund for Undergraduate Scholarly Experiences to A.C.G. along with the National Institutes of Health (R15MH113074), Nebraska EPSCoR First Award (OIA-1557417), Nebraska Research Initiative, and UNO start-up funds to RYW.

## Author Contributions

A.C.G. and R.Y.W. conceived the study, conducted statistical analyses, and wrote the manuscript. A.C.G. conducted the behavioral testing, brain extraction, gene expression quantification, and data collection.

## Figures Legends

**Figure S1.**
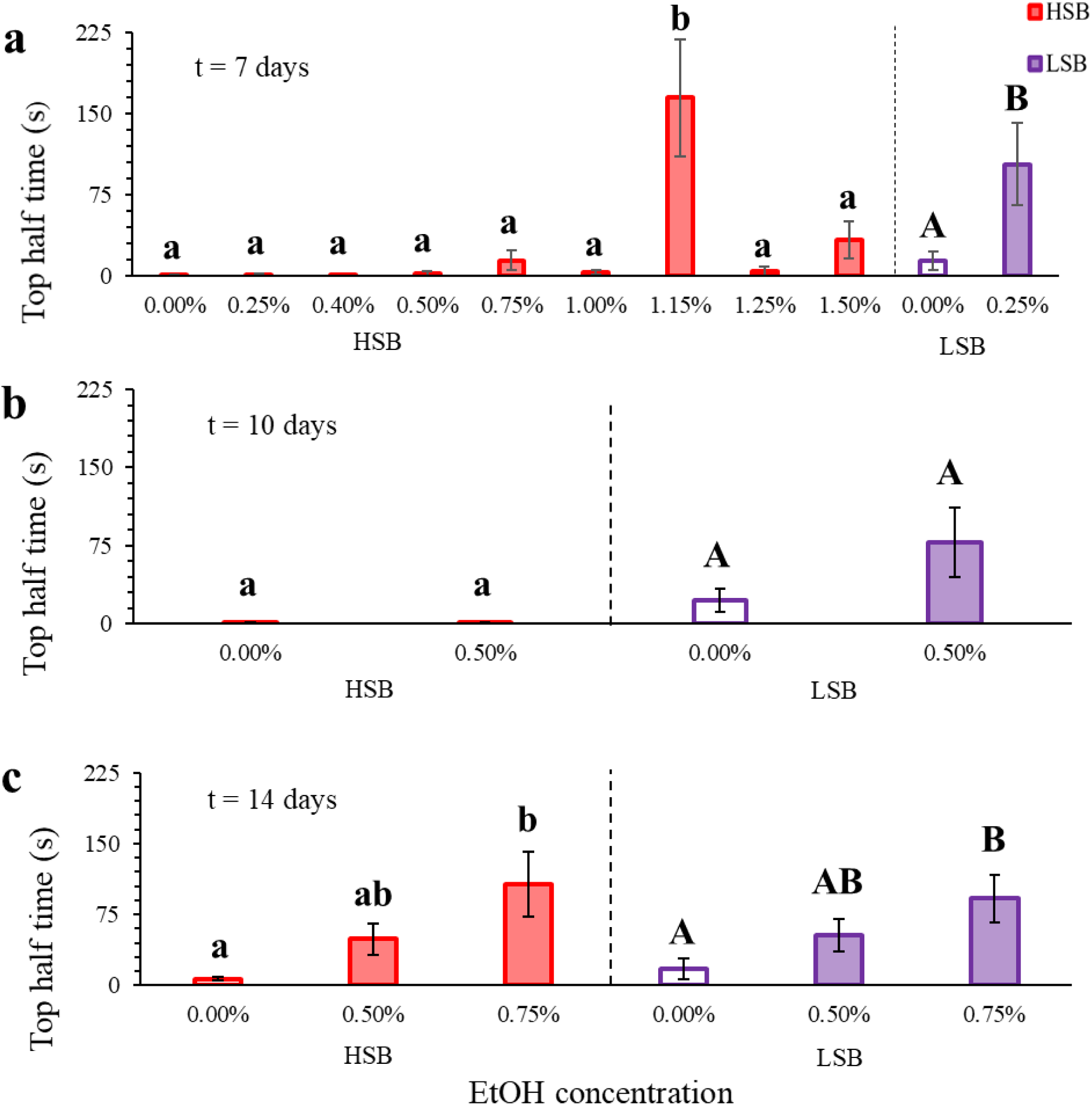
Dose response analysis of ethanol concentration on time spent in the top half of the tank during NTDT. Measured time spent in the top half of the tank after 7 days **(a)**, 10 days **(b)**, and 14 days **(c)** of each treatment. Control groups are represented by unfilled in bars, while ethanol-treated groups are represented by filled bars. HSB and LSB are red and purple, respectively. Data shown are mean ± 1 SEM. Individual differences within the HSB line are indicated by lower case letters, while differences within the LSB line are indicated by upper case letters.

**Table S1.** Summary of all generalized linear models ran for each of the ethanol treatment durations during the pilot dose-response study. Comparisons were made between treatment groups for each line at each duration. Abbreviations: HSB, high stationary behavior; LSB, low station behavior, 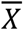, average; 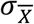, standard error; *χ*^*2*^, Chi-squared, U; *p*, p-value.

| Generalized Linear Model (GLZ) |  |  |  |  |  |  |  |
| --- | --- | --- | --- | --- | --- | --- | --- |
| Treatment Duration | Statistic | Time in Top (s) |  | Distance Traveled (cm) |  | Stationary Time (s) |  |
|  |  | HSB | LSB | HSB | LSB | HSB | LSB |
| 7-Day<br>$N_{\text{HSB}} = 94$<br>$N_{\text{LSB}} = 11$ | $\bar{X}$ | 21.566 | 108.497 | 688.475 | 1913.645 | 228.307 | 18.494 |
| | $\chi^2$ | 12.196 | 38.323 | 2.418 | 6.932 | 0.000 | 0.000 |
| | $U$ | 4.405 | 9.736 | 5.103 | 2.088 | 9.150 | 1.502 |
| | $p$ | 0.270 | $\leq 0.01$ | $\leq 0.05$ | 0.148 | $\leq 0.01$ | 0.220 |
| 10-Day<br>$N_{\text{HSB}} = 12$<br>$N_{\text{LSB}} = 11$ | $\bar{X}$ | 0.679 | 52.707 | 1362.251 | 2021.903 | 160.684 | 12.807 |
| | $\chi^2$ | $\leq 0.001$ | 0.123 | 0.003 | 0.020 | 0.000 | 0.000 |
| | $U$ | 0.122 | 2.640 | 41.798 | 0.010 | 87.957 | 0.998 |
| | $p$ | 0.727 | 0.104 | $\leq 0.001$ | 0.920 | $\leq 0.001$ | 0.318 |
| 14-Day<br>$N_{\text{HSB}} = 65$<br>$N_{\text{LSB}} = 46$ | $\bar{X}$ | 39.531 | 45.696 | 1728.181 | 1702.775 | 76.896 | 31.155 |
| | $\chi^2$ | 0.000 | 0.005 | 0.000 | 0.002 | 0.000 | 0.000 |
| | $U$ | 12.338 | 8.707 | 39.846 | 3.873 | 14.041 | 6.792 |
| | $p$ | $\leq 0.001$ | $\leq 0.01$ | $\leq 0.001$ | $\leq 0.05$ | $\leq 0.001$ | $\leq 0.01$ |

**Table S2.** Summary of Mann-Whitney U tests for the 7-day duration of ethanol treatment during the pilot dose-response study. Comparisons were made with control for all listed ethanol concentrations. Abbreviations: HSB, high stationary behavior; LSB, low station behavior, 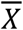, average; 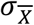, standard error; *U*, Mann-Whitney U; *p*, p-value; nt, not tested.

| 7-Day Ethanol Treatment Period |  |  |  |  |  |  |  |
| --- | --- | --- | --- | --- | --- | --- | --- |
| Ethanol Concentration | Statistic | Time in Top (s) |  | Distance Traveled (cm) |  | Stationary Time (s) |  |
|  |  | HSB | LSB | HSB | LSB | HSB | LSB |
| Control<br>$N_{\text{HSB}} = 34$<br>$N_{\text{LSB}} = 5$ | $\bar{X}$ | 0.840 | 31.417 | 453.697 | 1680.103 | 279.420 | 40.686 |
| | $\sigma_{\bar{X}}$ | 0.610 | 15.577 | 110.888 | 310.747 | 19.434 | 40.661 |
| 0.25% EtOH<br>$N_{\text{HSB}} = 10$<br>$N_{\text{LSB}} = 6$ | $\bar{X}$ | 0.671 | 172.730 | 931.620 | 2108.263 | 173.697 | $\leq 0.001$ |
| | $\sigma_{\bar{X}}$ | 0.638 | 43.467 | 331.941 | 153.014 | 43.019 | $\leq 0.001$ |
| | $U$ | 157.000 | 4.000 | 111.000 | 8.000 | 87.000 | 9.000 |
| | $p$ | 0.542 | 0.450 | 0.098 | 0.201 | $\leq 0.05$ | 0.104 |
| 0.4% EtOH<br>$N_{\text{HSB}} = 5$<br>$N_{\text{LSB}} = 0$ | $\bar{X}$ | 0.087 | nt | 631.633 | nt | 275.716 | nt |
| | $\sigma_{\bar{X}}$ | 0.055 | nt | 341.677 | nt | 36.151 | nt |
| | $U$ | 65.000 | nt | 64.000 | nt | 70.000 | nt |
| | $p$ | 0.181 | nt | 0.378 | nt | 0.529 | nt |
| 0.5% EtOH<br>$N_{\text{HSB}} = 6$<br>$N_{\text{LSB}} = 0$ | $\bar{X}$ | 2.314 | nt | 523.456 | nt | 269.770 | nt |
| | $\sigma_{\bar{X}}$ | 2.196 | nt | 209.965 | nt | 38.061 | nt |
| | $U$ | 81.000 | nt | 88.000 | nt | 87.000 | nt |
| | $p$ | 0.200 | nt | 0.596 | nt | 0.570 | nt |
| 0.75% EtOH<br>$N_{\text{HSB}} = 6$<br>$N_{\text{LSB}} = 0$ | $\bar{X}$ | 14.676 | nt | 1936.743 | nt | 45.139 | nt |
| | $\sigma_{\bar{X}}$ | 8.788 | nt | 279.091 | nt | 38.844 | nt |
| | $U$ | 43.000 | nt | 18.000 | nt | 14.500 | nt |
| | $p$ | $\leq 0.05$ | nt | $\leq 0.001$ | nt | $\leq 0.001$ | nt |
| 1% EtOH<br>$N_{\text{HSB}} = 12$<br>$N_{\text{LSB}} = 0$ | $\bar{X}$ | 3.896 | nt | 1062.793 | nt | 163.193 | nt |
| | $\sigma_{\bar{X}}$ | 2.171 | nt | 274.282 | nt | 41.211 | nt |
| | $U$ | 129.000 | nt | 110.000 | nt | 108.500 | nt |
| | $p$ | $\leq 0.01$ | nt | $\leq 0.05$ | nt | $\leq 0.05$ | nt |
| 1.15% EtOH<br>$N_{\text{HSB}} = 10$<br>$N_{\text{LSB}} = 0$ | $\bar{X}$ | 164.901 | nt | 462.236 | nt | 231.320 | nt |
| | $\sigma_{\bar{X}}$ | 54.315 | nt | 135.989 | nt | 40.515 | nt |
| | $U$ | 77.000 | nt | 145.000 | nt | 116.500 | nt |
| | $p$ | $\leq 0.01$ | nt | 0.484 | nt | 0.134 | nt |
| 1.25% EtOH<br>$N_{\text{HSB}} = 6$<br>$N_{\text{LSB}} = 0$ | $\bar{X}$ | 4.527 | nt | 486.142 | nt | 231.106 | nt |
| | $\sigma_{\bar{X}}$ | 4.527 | nt | 130.321 | nt | 45.216 | nt |
| | $U$ | 95.000 | nt | 65.000 | nt | 57.000 | nt |
| | $p$ | 0.810 | nt | 0.161 | nt | 0.088 | nt |
| 1.5% EtOH<br>$N_{\text{HSB}} = 5$<br>$N_{\text{LSB}} = 0$ | $\bar{X}$ | 33.326 | nt | 352.524 | nt | 259.476 | nt |
| | $\sigma_{\bar{X}}$ | 17.433 | nt | 103.970 | nt | 40.661 | nt |
| | $U$ | 38.000 | nt | 75.000 | nt | 61.000 | nt |
| | $p$ | $\leq 0.05$ | nt | 0.674 | nt | 0.313 | nt |

**Table S3.** Summary of Mann-Whitney U tests for the 10-day duration of ethanol treatment during the pilot dose-response study. Comparisons were made with control for all listed ethanol concentrations. Abbreviations: HSB, high stationary behavior; LSB, low station behavior, 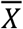, average; 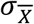, standard error; *U*, Mann-Whitney U; *p*, p-value.

| 10-Day Ethanol Treatment Period |  |  |  |  |  |  |  |
| --- | --- | --- | --- | --- | --- | --- | --- |
| Ethanol Concentration | Statistic | Time in Top (s) |  | Distance Traveled (cm) |  | Stationary Time (s) |  |
|  |  | HSB | LSB | HSB | LSB | HSB | LSB |
| Control<br>$N_{\text{HSB}} = 6$<br>$N_{\text{LSB}} = 5$ | $\bar{X}$ | 0.567 | 21.982 | 327.788 | 2038.106 | 304.745 | 0.267 |
| | $\sigma_{\bar{X}}$ | 0.486 | 11.234 | 129.727 | 215.263 | 29.910 | 0.027 |
| 0.5% EtOH<br>$N_{\text{HSB}} = 6$<br>$N_{\text{LSB}} = 6$ | $\bar{X}$ | 0.790 | 78.312 | 2396.714 | 2008.400 | 16.622 | 23.457 |
| | $\sigma_{\bar{X}}$ | 0.500 | 33.436 | 325.668 | 235.971 | 15.425 | 23.457 |
| | $U$ | 15.000 | 7.000 | 0.000 | 13.000 | 0.000 | 15.000 |
| | $p$ | 0.592 | 0.144 | $\leq 0.01$ | 0.715 | $\leq 0.01$ | 1.000 |

**Table S4.** Summary of Mann-Whitney U tests for the 14-day duration of ethanol treatment during the pilot dose-response study. Comparisons were made with control for all listed ethanol concentrations. Abbreviations: HSB, high stationary behavior; LSB, low station behavior, 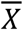, average; 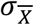, standard error; *U*, Mann-Whitney U; *p*, p-value.

| 14-Day Ethanol Treatment Period |  |  |  |  |  |  |  |
| --- | --- | --- | --- | --- | --- | --- | --- |
| Ethanol Concentration | Statistic | Time in Top (s) |  | Distance Traveled (cm) |  | Stationary Time (s) |  |
|  |  | HSB | LSB | HSB | LSB | HSB | LSB |
| Control<br>$N_{\text{HSB}} = 30$<br>$N_{\text{LSB}} = 23$ | $\bar{X}$ | 7.189 | 17.989 | 1067.222 | 1547.742 | 127.707 | 60.541 |
| | $\sigma_{\bar{X}}$ | 2.094 | 10.793 | 118.249 | 127.104 | 23.447 | 22.992 |
| 0.25% EtOH<br>$N_{\text{HSB}} = 24$<br>$N_{\text{LSB}} = 11$ | $\bar{X}$ | 49.040 | 53.281 | 2259.175 | 1759.733 | 45.476 | 3.695 |
| | $\sigma_{\bar{X}}$ | 16.214 | 17.264 | 193.773 | 1120.692 | 17.783 | 3.645 |
| | $U$ | 163.000 | 39.000 | 111.000 | 104.000 | 216.000 | 83.000 |
| | $p$ | $\leq 0.001$ | $\leq 0.001$ | $\leq 0.001$ | 0.408 | $\leq 0.01$ | 0.067 |
| 0.4% EtOH<br>$N_{\text{HSB}} = 11$<br>$N_{\text{LSB}} = 12$ | $\bar{X}$ | 106.989 | 91.850 | 2372.266 | 1947.711 | 6.871 | 0.003 |
| | $\sigma_{\bar{X}}$ | 34.732 | 25.571 | 247.109 | 154.496 | 4.090 | 0.003 |
| | $U$ | 18.000 | 49.500 | 33.000 | 78.000 | 70.000 | 78.000 |
| | $p$ | $\leq 0.001$ | $\leq 0.01$ | $\leq 0.001$ | $\leq 0.05$ | $\leq 0.01$ | $\leq 0.05$ |

**Table S5.**
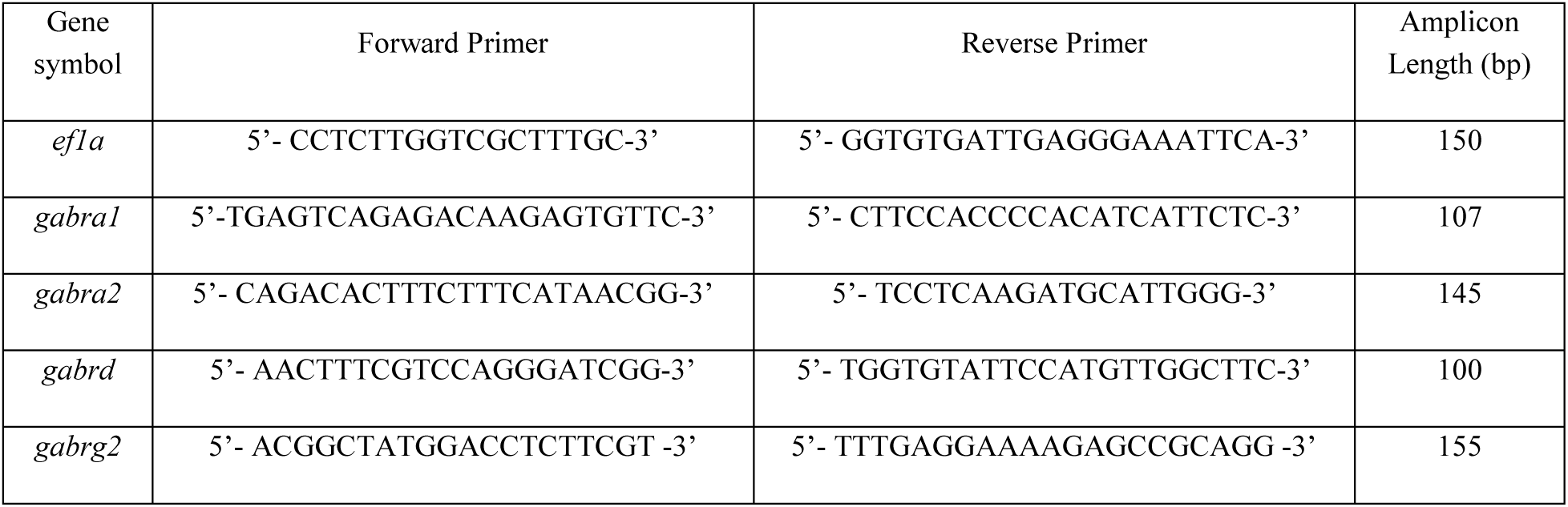
qRT-PCR primer characteristics.

